# Myofibrillar malformations that arise in mdx muscle fibers are driven by detyrosinated microtubules

**DOI:** 10.1101/2023.03.27.534405

**Authors:** Anicca Harriot, Tessa Altair-Morris, Camilo Venegas, Jacob Kallenbach, Kaylie Pinto, Humberto C. Joca, Marie-Jose Moutin, Guoli Shi, Jeanine Ursitti, Anna Grosberg, Christopher W. Ward

## Abstract

In Duchenne muscular dystrophy (DMD), alterations in the myofibrillar structure of skeletal muscle fibers that impair contractile function and increase injury susceptibility arise as a consequence of dystrophic pathology. In murine DMD (*mdx*), myofibrillar alterations are abundant in advanced pathology (>4 months), an age where we formerly established the densification of microtubules (MTs) post-translationally modified by detyrosination (deTyr-MTs) as a negative disease modifier. Given the essential role of MTs in myofibrillar growth, maintenance, and repair, we examined the increased abundance of deTyr-MTs as a potential mechanism for these myofibrillar alterations. Here we find increased levels of deTyr-MTs as an early event in dystrophic pathology (4 weeks) with no evidence of myofibrillar alterations. At 16 weeks, we show the level of deTyr-MTs is significantly increased and co-localized to areas of myofibrillar malformation. Profiling the enzyme complexes responsible for deTyr-tubulin, we identify vasohibin 2 (VASH2) and small vasohibin binding protein (SVBP) significantly elevated in the *mdx* muscle at 4 wks. We then use the genetic increase in VASH2/SVBP expression in 4 wk wild-type mice and find densified deTyr-MTs that co-segregate with myofibrillar malformations similar to those in the 16 wk *mdx*. Given that no changes were identified in fibers expressing EGFP as a control, we conclude that disease dependent densification of deTyr-MTs underscores the altered myofibrillar structure in dystrophic skeletal muscle fibers.

## 1. Introduction

Skeletal muscle fibers exhibit highly ordered myofibrillar structure which is essential for efficient force generation within the muscle fiber. Myofibrils (1µm diameter) are composed of individual contractile units (i.e., sarcomeres, ∼2 µm in length) arranged in series to span the length of the muscle fiber (500µm to a few cm). The number of parallel packed myofibrils within the muscle fiber governs the contractile force. Until recently, myofibrils were thought to be independent units co-registered through protein links between their Z-line sarcomere boundaries. However, new evidence of sarcomeres branching between registered myofibrils^1^ has redefined these structures as a continuous myofibrillar matrix that facilitates the highly coordinated, unilateral contraction of the muscle fiber.

In contrast to the registered myofibrillar matrix seen in heathy skeletal muscle, are myopathies such as Duchenne muscular dystrophy (DMD) where myofibrils become misaligned and torturous resulting in misorientation of force vectors and dyssynchronous activation of sarcomeres^2–5^. These changes result in significant reductions in isometric force and velocity of contraction as well as increased shear-stress that predisposes damage at these locations ^6–8^. While misalignment of the sarcomeres is now established as pathognomonic in DMD, the mechanisms that predispose their occurrence are unknown.

The cytoskeleton is a dynamic structural and signaling scaffold of microtubule (MT), actin and intermediate filaments (IF) that is essential for the intracellular trafficking, maintenance of cellular architecture, and positioning of organelles in all cells. Microtubules garnered early attention in DMD muscle as a cytoskeletal element that was disorganized early in disease and became densified with disease progression^9–11^. The discovery of dystrophin as an MT binding partner has resolved the mechanisms that initiate the disorganized MT structure in DMD^11^ and inspired many groups to determine how these MT alterations may impact dystrophic pathology.

The structure and function of MTs are regulated by post-translational modifications (PTM) to their tubulin monomers. Detyrosination (deTyr), the reversible enzymatic removal of α-tubulin’s COOH-terminal tyrosine, promotes the interaction of MTs with binding partners^12–15^. Work by our group has identified that MTs enriched in deTyr-tubulin (deTyr-MTs) regulate the stiffness of the muscle fiber cytoskeleton and thus the activation of NADPH Oxidase 2 (Nox2) dependent reactive oxygen species (ROS) and calcium (Ca^2+^) signals by mechanotransduction^16, 17^. In the murine model of DMD (i.e., *mdx*), the densification of deTyr enriched MTs is a consequence of disease pathology that increases the passive mechanics of muscle fibers. Together with the increased expression of Nox2 proteins, these changes drive the excess mechanotransduction elicited Nox2- ROS and Ca^2+^ signals linked to dystrophic pathology^16,17,18^.

Our lab’s previous work on dysregulated MT mechanotransduction in murine DMD (*mdx*) focused on murine models between 3-9 months, when pathology is entrenched yet progression is evident^16, 17^. Within this timeframe of disease our group and others have previously identified and profiled the increased occurrence of muscle fibers with gross structural malformations (i.e., splitting, branching)^2, 3, 7, 19, 20^ that increase the susceptibility to contractile damage in DMD. Here we were intrigued by work suggesting that these gross alterations in muscle fiber structure arose from structural changes in the myofibrils^3^. Informed by our observation of the MT densification often occurring in discrete areas in *mdx* muscle fibers^16^ ^17^, and evidence that microtubules are essential for myofibrillar growth, maintenance, and repair^21–24^, we hypothesized a link between the disease altered MTs and the occurrence of myofibrillar malformations in DMD.

In the present study, we focused early in disease pathology to bias our capture of the mechanisms that underlie the development of myofibrillar malformations. In muscle fibers from young mice (4 wks) we identify a small but significant increase in deTyr-MTs in *mdx,* yet no evidence of myofibrillar malformation above WT. Profiling muscle fibers at 16 wks, we find the level of deTyr-tubulin increases disproportionally in the *mdx* where it occurs largely in bundles of MTs that co-localize with areas of myofibrillar malformation.

Microtubules participate in the highly orchestrated growth, maintenance, and repair of myofibrillar structure^24^. In fact, an increased level of deTyr enriched MTs is an early and critical event in sarcomerogenesis^25, 26^. Profiling the enzyme complexes responsible for deTyr-tubulin we found vasohibin 2 (VASH2) significantly elevated in the *mdx* muscle at 4 and 16 wks. To determine the consequences of elevated VASH2 activity on myofibrillar structure we overexpressed VASH2 in muscles of 4wk wild-type mice. We show that VASH2 overexpression modeled the densification of deTyr-MTs seen in the 16 wk *mdx*. Furthermore, we demonstrate that these deTyr-MTs co- segregate with myofibrillar malformations comparable to those found in 16wk *mdx*. We conclude that disease altered microtubules are an early event in dystrophic pathology that predisposes the altered myofibrillar structure in dystrophic skeletal muscle fibers.

## 2. Methods

### 2.1 Animal Use

All mice were obtained from Jackson Laboratories (Bar Harbor, MA) and studied with procedures approved by the University of Maryland, Baltimore Institutional Animal Care and Use Committee (IACUC).

#### 2.1.1 Electroporation

Anesthetized mice (2% isoflurane) were injected with 25 µL of 1 mg/mL of hyaluronidase (Sigma-Aldrich) subcutaneously into the sterilized food pad of both hindlimbs. After 1 h, one footpad was injected with plasmid cDNA (20 µL at 1 µg/µL) containing a bicistronic construct of VASH2-GFP/SVBP (kind gift from Marie Jo Moutin^27^) with the contralateral footpad receiving a plasmid cDNA containing EGFP as a control (1 µg/µL). The plasmid cDNA was then delivered to the flexor digitorum brevis (FDB) muscle by electroporation through sterile electrodes placed subcutaneously at the proximal and distal ends of the FDB. The pulse protocol consisted of 30 pulses of 150 V for 20 ms duration at a frequency of 1 Hz. Mice were humanely euthanized and FDBs harvested after 5 days.

#### 2.1.2 Muscle Fiber Preparation

FDB muscles were harvested bilaterally in sterile mouse ringer and maintained overnight in DMEM (supplier) supplemented with collagenase A (Roche, 0.2 mg/ml) and 1% Penicillin-Streptomycin in a CO_2_ incubator (37°C, 5% CO_2_). Following gentle trituration to yield single FDB fibers, the cells were washed once in DMEM supplemented with 10% fetal bovine serum then washed twice in physiological Ringer solution with 1mM EGTA (pH, 7.4). Fibers were then either maintained in Ringer solution at room-temperature for live cell imaging or fixed in 4% PFA with 5mM EGTA for 20 minutes at room temperature, washed twice in PBS then stored in PBS with 0.4% sodium azide until used.

#### 2.1.3 In-vivo contractile function

Contractile performance and injury susceptibility were tested *in vivo* as described previously^16, 17^. Anesthetized mice (2-3% isoflurane) were placed in supine position on the temperature maintained (Deltaphase Isothermal Pad, Braintree Scientific) platform of an Aurora 3100 with the knee stabilized and foot affixed on the footplate of the torque transducer. The plantar flexor muscle group (gastrocnemius, soleus) was activated by percutaneous stimulation. The force frequency relationship was evaluated with 500msec trains of square pulses (0.1 ms) between 1 and 150Hz. The susceptibility to contraction force-loss was evaluated with 25 eccentric (i.e., lengthening) contractions.

#### 2.1.4 Mechanical properties

FDBs maintained in Ringer were placed on a glass-bottom dish coated with ECM (E6909; Sigma-Aldrich). The near membrane mechanical properties of the FDB were quantified with a Chiaro nano-indenter (Optics11) using a cantilever (0.044 N/m stiffness) with round probe (3-µm radius). Indentation (1uM) profiles at speeds from 0.5 to 25 µm/s were analyzed with a Hertzian contact model to calculate the Young’s modulus (i.e., stiffness) of the FDB.

### 2.2 Western Blotting

Western blots were conducted as previously described^17^. Briefly, 20μg of clarified muscle was processed via SDS-PAGE, transferred to nitrocellulose membranes, and washed with 5% milk solution in PBS before blocking for 1h in the same solution. The membrane was probed overnight for β-tubulin (T4026, Sigma-Aldrich), deTyr-tubulin (31-1335-00, RevMAb Biosciences), acetylated tubulin (T7451, clone 6-11B-1; Sigma-Aldrich), and gp91phox (Abcam; ab129068). Next membranes were washed twice in 5% milk solution, incubated with the appropriate secondary antibody (1:10,000) at room temperature for 1h, and washed with 0.5% Tween solution in PBS twice for 10min. Blots were imaged and analyzed using the LICOR Odyssey CL-x system.

### 2.3 RT-qPCR

Gastrocnemius muscles were collected from *mdx* and WT mice and snap-frozen in isopentane cooled on dry ice. Tissues were later powdered and homogenized in TRI-reagent (Zymo Research). Phase separation was performed using 0.2mL of chloroform per 1mL of TRI-reagent, with samples shaken vigorously for 2min then centrifuged at 12,000 x g for 10min at 4°C. To precipitate RNA from the aqueous phase, 0.5mL isopropyl alcohol per 1 ml TRI-reagent used for lysis was added and incubated at room temperature for 10 minutes before centrifuging for 10 minutes at the aforementioned settings. The resulting RNA pellet was washed with 75% ethanol, centrifuged for 5 minutes at 7,500 x g at 4°C, then dissolved in 30µL DNase-RNase free water at which point RNA concentration was measured using a spectrophotometer. 2.5 µg samples of RNA were reverse transcribed using the SuperScript IV First-Strand Synthesis System (Invitrogen), following manufacturer protocol. Gene expression was measured via RT-qPCR using PowerUp SYBR Green Master Mix (Applied Biosystems) on the QuantStudio3 Real-time PCR System (Applied Biosystems by Thermo Fisher Scientific) and QuantStudio Design and Analysis software v1.5.2 (Applied Biosystems). Primers are listed in Supplementary Table 1.

### 2.4 Immunofluorescence and automated imaging

Fixed FDB fibers were blocked for 2h at room temperature in Superblock™ Blocking Buffer in PBS (Thermo Scientific) with 0.04% saponin. Fibers were incubated in an Eppendorf tube with primary antibodies to detect microtubule structure (beta-tubulin; T4026, Sigma-Aldrich) and the population of microtubule tubulin modified by detyrosination (deTyr-tubulin; 31-1335-00, clone RM444, RevMAb Biosciences USA, Inc.). Sarcomeric actin was decorated with phalloidin conjugated to Alexa Fluor 633 (A22284, Invitrogen) to visualize myofibrillar structure. Primary Antibodies and phalloidin were used overnight at 4**°**C. The following day FDB fibers were incubated with the appropriate secondary antibodies (diluted in PBS containing 0.04% saponin and 0.1% sodium azide) for 2 h at room temperature, washed three times in PBS, then mounted onto slides with ProLong Gold + Dapi mountant (Invitrogen).

Fixed FDB fibers were imaged on an inverted Nikon C2+ confocal fluorescence system using an automated protocol developed in NIS Elements AR JOBS. Single fibers were identified by their actin labeling (i.e., phalloidin 633) from a full-slide tile scanned image (10x air obj.). Fibers without evidence of bends or hypercontraction under visual inspection (30-50 fibers per slide) were logged as regions-of-interest (ROI). Each identified fiber ROI was imaged using an automated routine that identified the muscle fiber surface and collected a full thickness z-stack (0.5 µm steps; 4 frame average) at 40x (1.4 N.A. Plan Apo air obj.) and 1.3 Airy units which yielded 0.31 μm/pixel resolution.

#### 2.3.1 Automated microtubule structural analysis

The properties of the microtubule network were quantified in NIS Elements AR General Analysis 3. Briefly, an inverse binary mask of the phalloidin label (Cy5 channel) identified areas of myofibrillar structure (black) and areas of myofibrillar gaps (white) linked to the regions of altered continuity. Within each z-stack image the density of deTyr-tubulin (binarized deTyr- tubulin) was determined within the areas of myofibrillar structure and continuity gaps and normalized to the measured area. The area around the nuclei was masked to exclude any microtubule alterations around the nuclei as a confounding factor. For global density measures, the total deTyr-tubulin stain for each z-slice was normalized to the fiber area (as determined by the phalloidin label).

## 3. Results

### 3.1 Young *mdx* mice exhibit functional deficits and microtubule alterations

Our lab’s previous work on dysregulated MT mechanotransduction and gross structural alterations in *mdx* was in mice 3-9 months of age when pathology is well-established and still progressing. In this study, we examined wild-type (C57BL.10/J) and *mdx* (C57BL.10/J *mdx*) mice at 4 and 16 wks of age to elucidate the mechanisms that underlie the development of myofibrillar malformations. Our initial experiments sought to establish the functional status of the muscle at these ages. Evaluating *in vivo* plantar flexor function, we confirmed deficits in maximal isometric force in the *mdx* at both 4 and 16 wks (**Fig 1A**). Measuring the weight of the gastrocnemius muscle we identified no differences between genotypes at 4 wks yet a significant increase in the mass of the *mdx* gastrocnemius at 16 wks, a finding consistent with the pseudohypertrophy reported at this age (**Fig 1B**). Calculating the specific force (i.e., force normalized to mass) revealed no difference between genotypes at 4 wks yet a significant drop in the specific force of the *mdx* was observed at 16 wks (**Fig 1C**). Finally, evaluating isometric force loss following 20 *in vivo* eccentric contractions we again found no significant deficit between genotypes at 4 wks; this however, progressed to a significant decrease in *mdx* at 16 wks (**Fig 1D**). Taken together, we identified an acceleration in functional deficits after 4 wks of age in the *mdx*.

**Fig 1.**
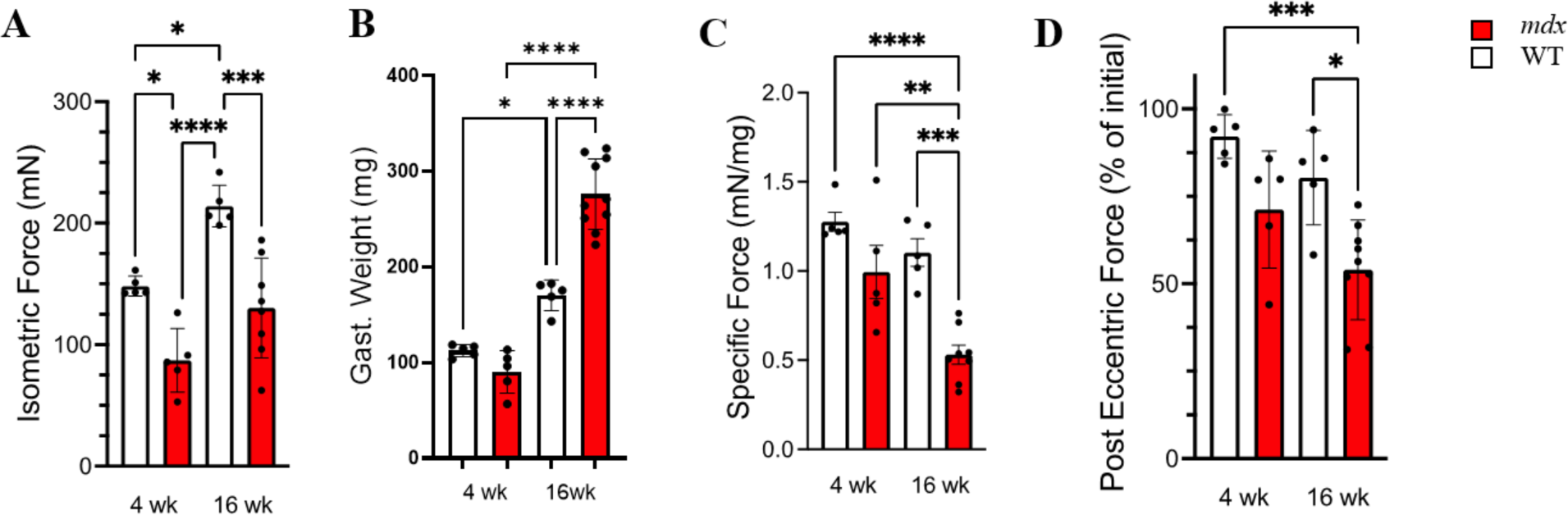
Force production decreases with age in *mdx* while injury susceptibility increases. **(A)** Isometric force produced at 150Hz in wild-type (white) and *mdx* (red) gastrocnemius muscles at 4 and 16 wks, respectively. **(B)** Weights of the gastrocnemius muscles from WT and *mdx* mice used to normalize force production to measure **(C)** specific force production in age-matched WT and *mdx* mice at 150 Hz. **(D)** Percent force production in gastrocnemius muscles from 4 and 16 wk WT and *mdx* mice after 25 eccentric contractions. *Mdx* mice experience increased force loss when compared to wild-type. Values are means ± SEM. All analysis are one-way ANOVA with Šídák’s multiple comparisons test (* P < 0.05; ** P < 0.01; ***P<0.001; ****P <0.0001)

Our group, and others^28–31^, have implicated disease dependent MT alterations as negative disease modifiers in adult *mdx* mice with advanced pathology. Western blot profiling of gastrocnemius muscle from 4 wk *mdx* mice finds no significant alteration in the expression of tubulin protein, its modification by acetylation (acetyl), nor the abundance of gp91phox, the catalytic subunit of the reactive oxygen species (ROS) producing enzyme, Nox2. While not statistically significant, detyrosination (deTyr) shows a marked increase at this age (**Fig 2**). As disease progresses from 4 to 16 wks we identify an increased abundance of gp91phox, increased expression of tubulin protein as well as its modification by detyrosination (deTyr) and acetylation. Our published works have linked these tubulin changes to the increased myofiber passive mechanics (i.e., stiffness) and together with gp91phox, to the excess MT dependent mechanotransduction activation of Nox2-ROS and Ca^2+^ signals that drive disease pathology in DMD.

**Fig 2.**
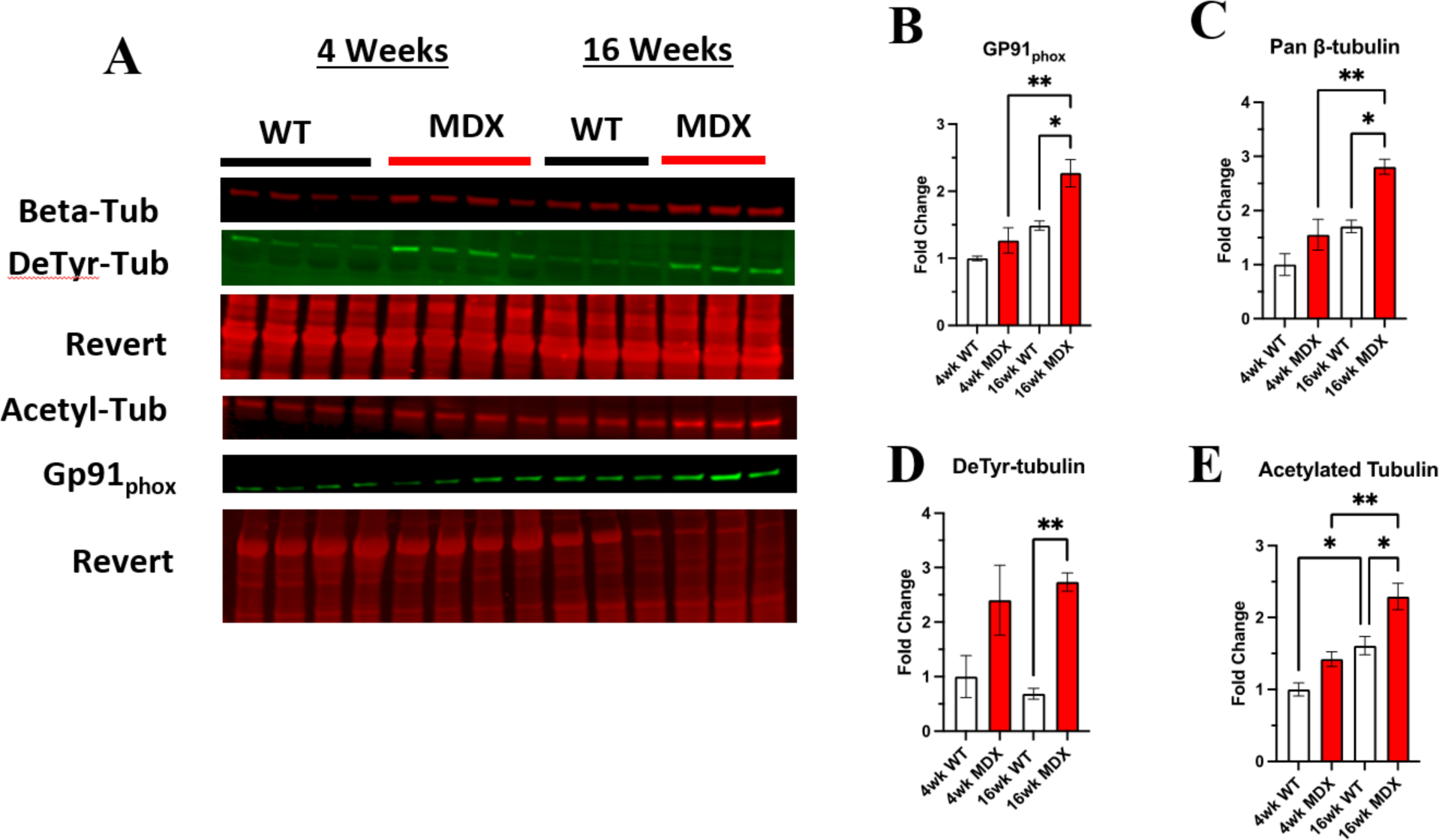
Detyrosination increases with dystrophic disease progression. **(A)** Western blots of the gastrocnemius muscle from wild-type and *mdx* mice at 4 wks (n=4) and 16 wks (n=3) **(B)** increased tubulin abundance and post-translational modification by **(C)** detyrosination and **(D)** acetylation. **(E)** We also see a significant increase in the NOX2 subunit GP91_phox_. Values are means ± SEM. All analysis are one-way ANOVA with Šídák’s multiple comparisons test (* P < 0.05; ** P < 0.01)

### 3.2 *mdx* Mice Exhibit Malformed Myofibrillar Structure

Alterations in myofibrillar structure are a consequence of deficient myofibrillar repair following acute muscle damage^32^ or disease pathology^2, 3, 19^ that predisposes the occurrence of gross malformations in muscle structure (i.e., split and branched fibers)^2^. Our group previously reported a low percentage of grossly malformed (i.e., bifurcated, split, etc.) skeletal muscle fibers in 6-9 wk *mdx*, with less than 10% abnormal fibers found in the flexor digitorum brevis (FDB)^33, 34^. These observations were made by visual inspection in brightfield where we found > 90% of the FDB muscle fibers having no detectable abnormalities. In the current study, we show that when labeled for myofibrillar structure (i.e., phalloidin labeled actin), and imaged with confocal microscopy, a more significant number of muscle fibers with abnormalities in myofibrillar structure becomes apparent (**Fig 3**).

**Fig 3.**
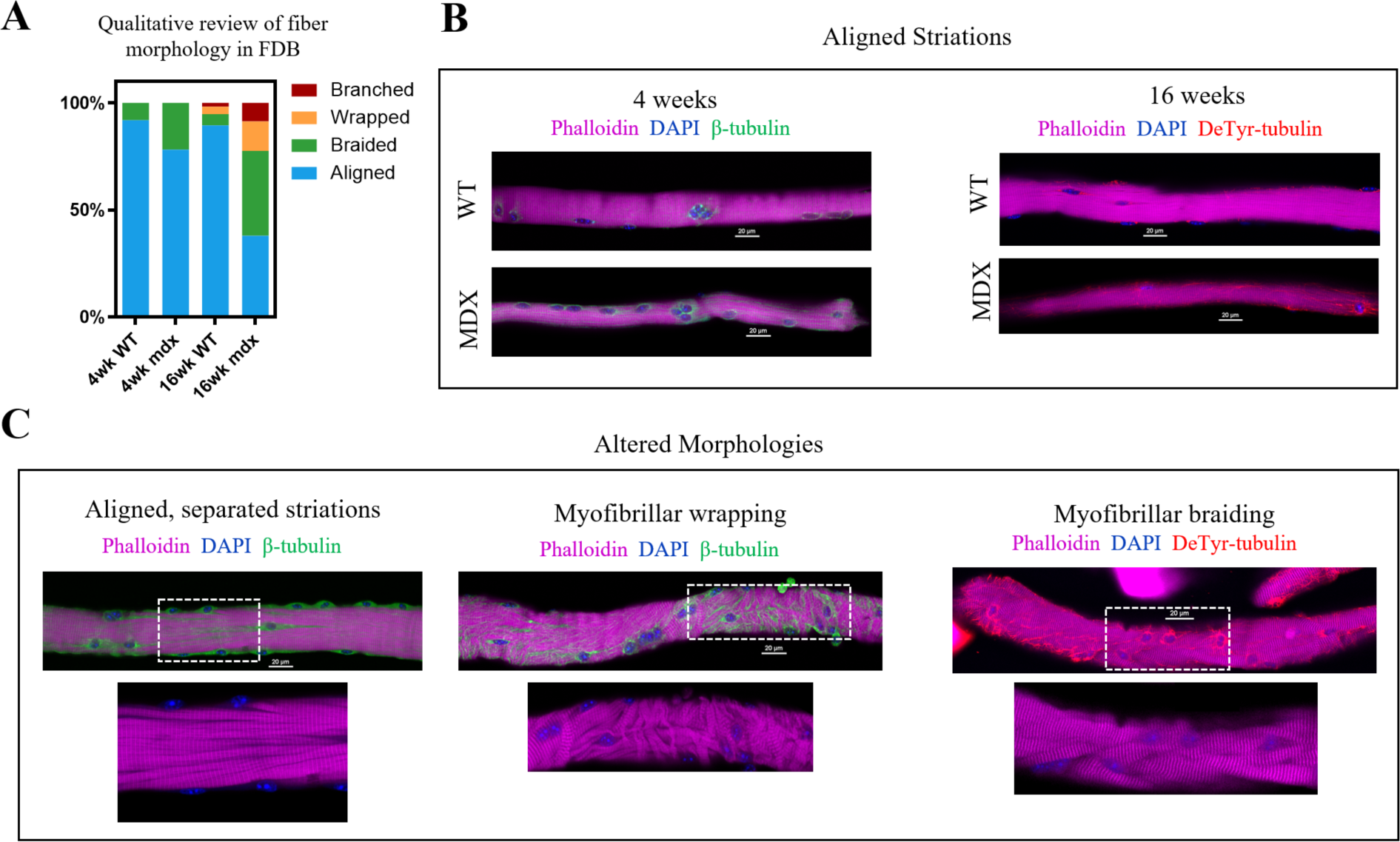
Qualitative survey of fiber morphology. **(A)** The percent distribution of each morphology was determined based on a survey of approx. of 150 fibers (n = 5, ∼30 fibers/animal) from each condition. **(B)** Representative images showing aligned striations at 4 and 16 wks with canonical rectilinear MT structure. MT bundling is apparent in *mdx* as early as 4wks. Detyrosinated tubulin appears increased in 16 wk *mdx*. **(C)** Fibers from 16 wk *mdx* representative of altered morphologies, showing areas of MT bundling appearing coincident with alterations in striation continuity, in fibers with myofibrillar malformation as well as in fibers with otherwise aligned striations.

Using an automated confocal strategy, we imaged single isolated FDB myofibers from WT and *mdx* at 4 wks and 16 wks. Our qualitative visual inspection identified four distinct morphologies of myofibrils within the muscle fiber: (1) canonical aligned striations; (2) evidence of “braided” myofibril structure with misregistration; (3) misalignment characterized by myofibrils wrapping around the peripheral myofibrils; (4) fibers exhibiting gross malformations (i.e., branches, splits) as previously described^33^ (**Fig 3A-C**). In the 4 wk *mdx*, myofibers with evidence of braided myofibrils make up 22% of the total FDB population. By 16 wks, only 38% of myofibers in the *mdx* FDB exhibit canonical aligned striations; 39% of fibers have braided myofibrils, 14% have myofibrillar wrapping, and 9% of muscle fibers are branched (**Fig 3C**). Although muscle fibers with altered myofibrillar morphologies were identified in the wild-type, these comprised less than 11% of the total FDB population.

While fibers with canonical aligned striations were in the majority in both genotypes, a significant number of otherwise normal *mdx* fibers presented with separations between myofibrils, marked by bundles of microtubules (**Fig 3C)**. In fact, these myofibrillar separations were evident in a majority of 16 wk *mdx* muscle fibers and were observed in some wild-type FDBs albeit at a markedly reduced occurrence. These initial qualitative observations provided the basis for our adopting quantitative methods. Informed by our past observations of the MT densification often occurring in discrete areas in *mdx* muscle fibers (Khairallah [Fig 2], Kerr [Fig 1]), and new qualitative evidence that this MT densification is coincident with altered myofibrillar structure, we posited a link between the disease altered MTs and the occurrence of myofibrillar malformations in DMD.

### 3.3 The Continuity of Z-line Striations as a Metric of Myofibrillar Structure

We next quantified the continuity of myofibrillar Z-line striations as a metric of myofibrillar structure. The quantitative assessment of Z-line striations in each image was performed using a custom MATLAB routine established by Morris et al.^35^ for profiling myofibrillar structure in developing cardiomyocytes and skeletal myotubes and adapted here for mature skeletal muscle fibers.

In brief, multi-channel Nikon confocal fluorescence images were converted into RGB TIFFs and the Cy5 channel containing phalloidin-633 for actin was output to a new image stack. To decrease processing time, each image was cropped to the fiber of interest and rotated to align the myofiber long-axis to the x-axis of the frame. The maximal myofiber boundary was identified with an Otsu’s threshold^36^ of the maximum intensity projection (max-IP) of the z-stack (**Fig 4A**). In the resulting binary image, the myofiber area was calculated and orientation axis determined using the least mean square orientation estimation algorithm. Subsequently, each z-slice was binarized based on Otsu’s threshold and compared to the binarized maximum intensity projection to determine the ratio of “true” pixels in the z-slice of interest compared to the max-IP (**Fig 4B-D**). Z-slice ratios above a user defined threshold (nthresh = 0.6) were selected for analysis (**Fig 4E**).

**Fig 4.**
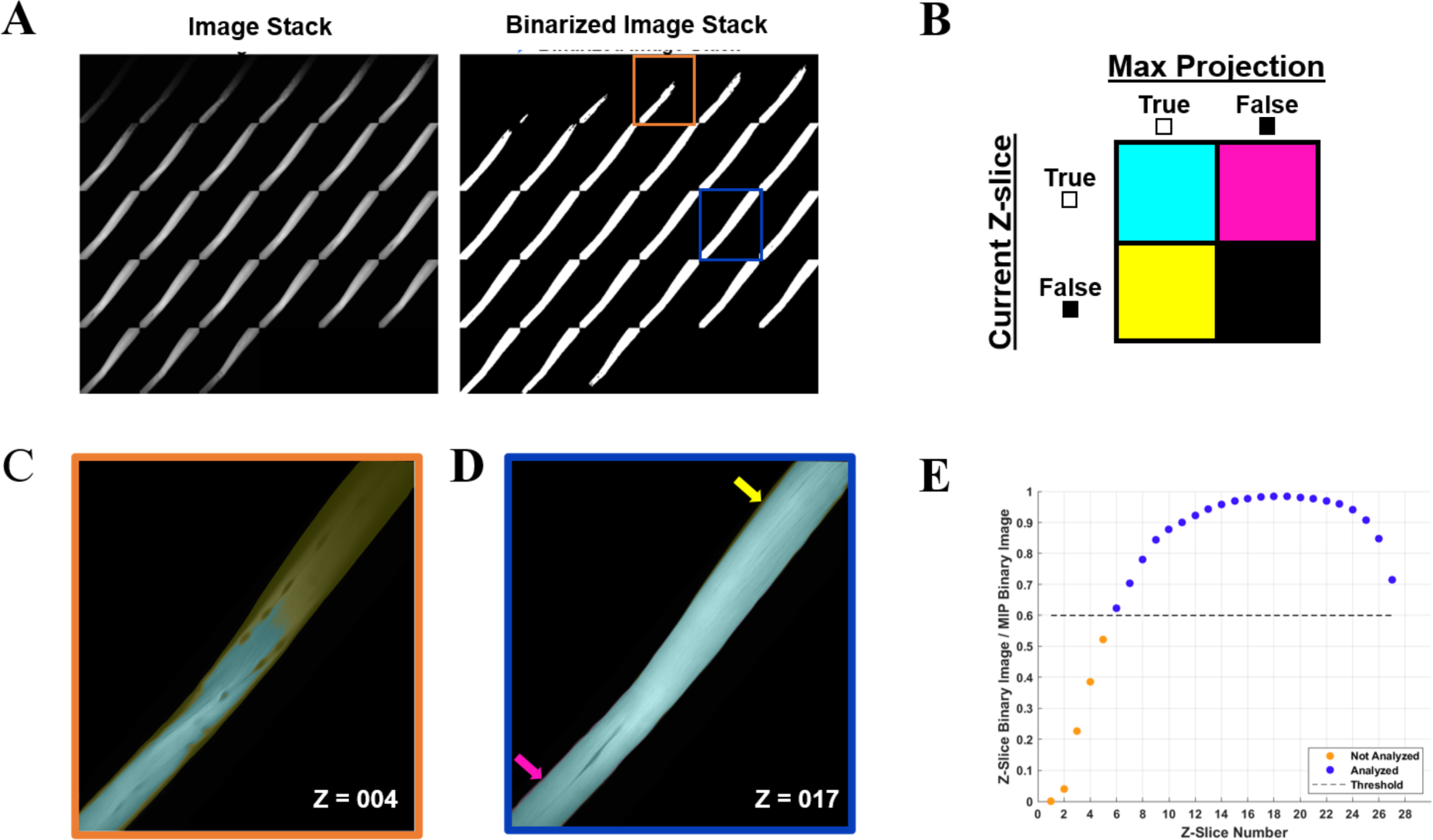
Automated image selection for myofiber analysis. **(A)** Each z-slice is binarized and compared with the binarized maximum intensity projection (max-IP). **(B)** Within the current z-slice pixels with sufficient intensity for analysis are logged as true. The total number of true pixels in the current z-slice is compared as a ratio between true pixels in the max-IP. **(C)** A representative selection shows the fiber area of 4^th^ z-slice (outlined in orange) was less than 60% of the max-IP area whereas **(D)** the fiber area of 17^th^ z-slice exceeded the 60% threshold. **(E)** A representative plot of the z-slices to be included in analysis based on the user defined threshold (nthresh = 0.6).

For each analyzed z-stack image, the minor axis length (i.e., width) of the myofiber was determined for every 20 pixels along the long axis of the muscle fiber (**Fig 5A**). Subsequently the phalloidin labeled Z-line structure was detected via the ZlineDetection algorithm developed by Morris et al.^37^. The continuity of each Z-line was determined by its length divided by the nearest minor axis length and plotted with color code **(Fig 5B**) with a continuous Z-line spanning the muscle fiber perpendicular axis yielding a measure of 1 (red), with interruptions in the continuity yielding lower values and cooler colors. The output for each fiber includes a mean striation continuity score for each z-slice as well as a boxplot for the entire z-stack (**Fig 5C**). The median striation length for the entire muscle fiber z-stack is also reported as the continuity score.

**Fig 5.**
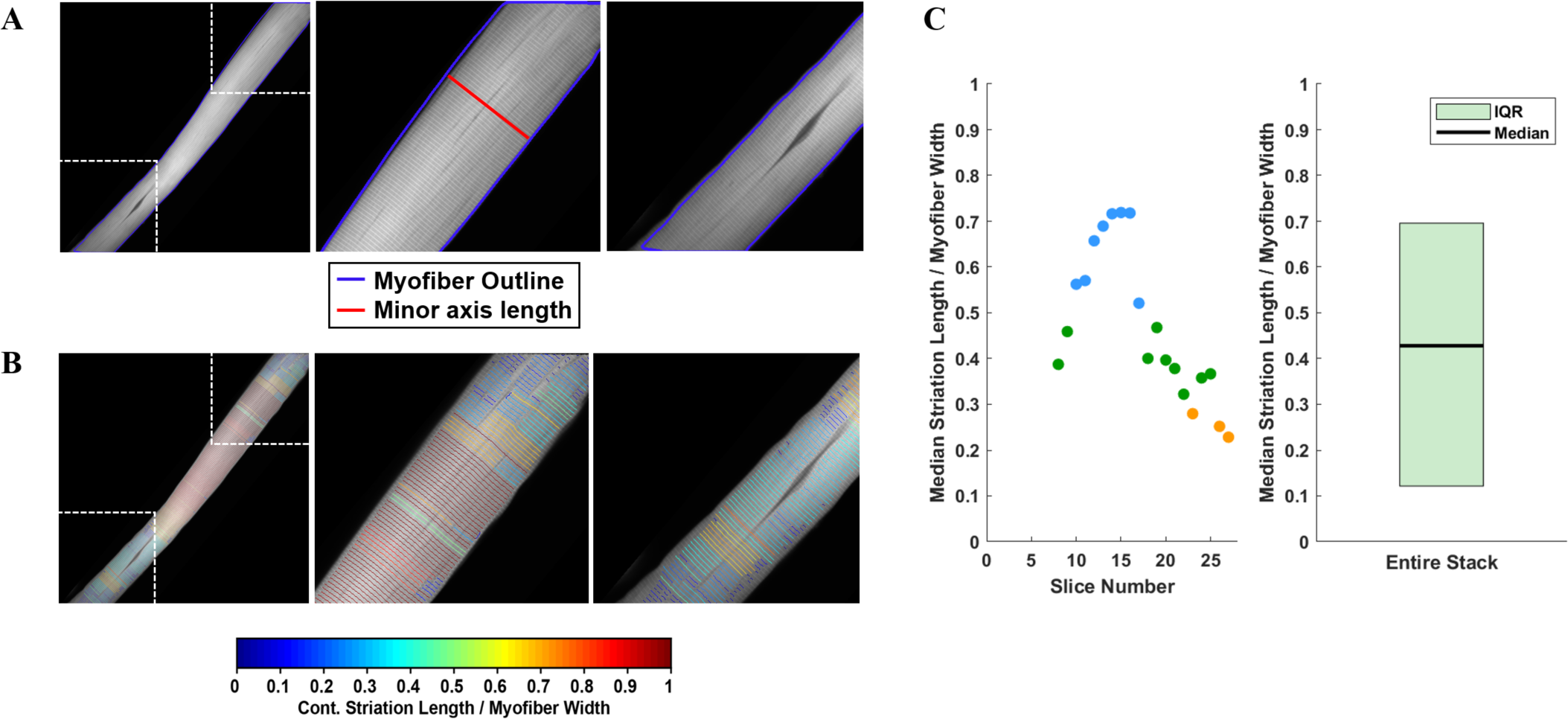
Striation continuity quantification. **(A)** Representative z-slice of phalloidin channel of WT FDB shown in Fig. 2. Insets (right) showing Z-line structure. **(B)** Automated Z-line detection output from ZlineDetection . Insets (right) showing the heterogeneity of Z-line continuity in WT FDB at minor separations between myofibrils and the nucleus. **(C)** Quantification of the striation length as compared to myofiber width within each z-slice (left) and the corresponding box plot for the entire z-stack (right).

### 3.4 Z-line continuity decreases with dystrophic progression and predicts altered myofibrillar structure

We show that striation continuity effectively identifies the minor interruptions in the myofibrillar structure seen in WT muscle fibers as well as the more significant disruptions in the *mdx* (**Fig 6A-F**). We demonstrate continuity scores < 0.4 in fibers displaying areas of myofibrillar braiding (**Fig 6B,D**), while scores decrease to < 0.2 in fibers with significant myofibrillar wrapping (**Fig 6C,F**).

**Fig 6.**
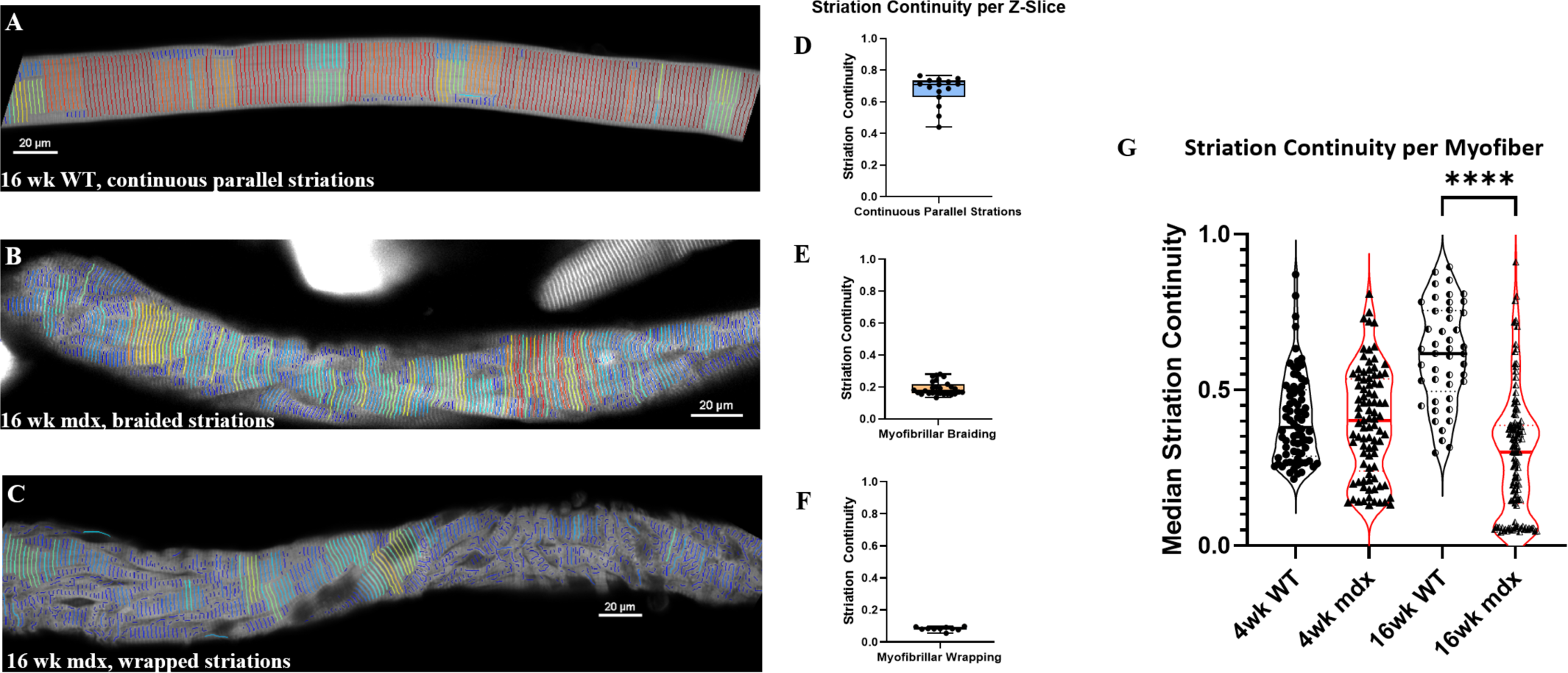
Median Striation Continuity scores decrease with altered morphology. Striation detection for (A) the full 16wk wild-type fiber depicted in Figure 1, (B) the 16wk mdx fiber with braided myofibrils, and (C) the 16wk mdx fiber with wrapped myofibrils. Their respective continuity scores per z-slice are illustrated with min, max, median scores displayed in boxplots (E-F). (G) Average striation continuity scores for each fiber within each experimental group (n=50- 125 fibers per condition). One-way ANOVA revealed significant difference between groups with Šídák’s multiple comparisons test elucidating a significant difference between 16wk WT and MDX (*F*(3, 333) = 38.36, *p* < 0.0001).

Using the automated imaging strategy and ZlineDetection to quantitate Z-line continuity, we screened FDB myofibers from WT and *mdx* mice at 4 wks and 16 wks of age. At 4 wks we find no significant difference in the mean striation continuity of the entire fiber between *mdx* and WT (**Fig 6G**), the 4-wk *mdx* do achieve lower minimum continuity scores than the 4-wk wild-type. Because severely misaligned myofibrils are a rare event at 4 wks, the reduced continuity score is manifest from microtubule bundles between myofibrils causing separations as previously described (**Fig 3A**).

By 16 wks, the differences between morphology in *mdx* and wild-type become more pronounced. There is an increase in the average continuity score in wild-type, while the continuity of the *mdx* decreases as altered morphologies emerge at a greater frequency (**Fig 6G**). This increase in the proportion of separations between myofibrils in the mdx between 4 and 16 wks suggests a progression of myofibrillar alterations with disease progression.

### 3.5 Increased tubulin detyrosination occurs commensurate with myofibrillar malformations

Examining FDB myofibers labeled for deTyr-tubulin and actin we find a significant increase in the density of deTyr-MTs in the *mdx* at 4 wks that progresses at 16 wks (**Fig 7A-B**). Revisiting our previous observation of bundled microtubules coinciding with myofibrillar separations, we quantified the density of deTyr-tubulin within the regions of myofibrillar separation versus the density in areas with otherwise normal myofibrillar connectivity (**Fig 7C-E**). At 4 wks we find no difference in the density of deTyr -MTs between these areas in WT, but in the *mdx* we find a significant increase in deTyr-MTs only in the regions of myofibrillar separation.

**Fig 7.**
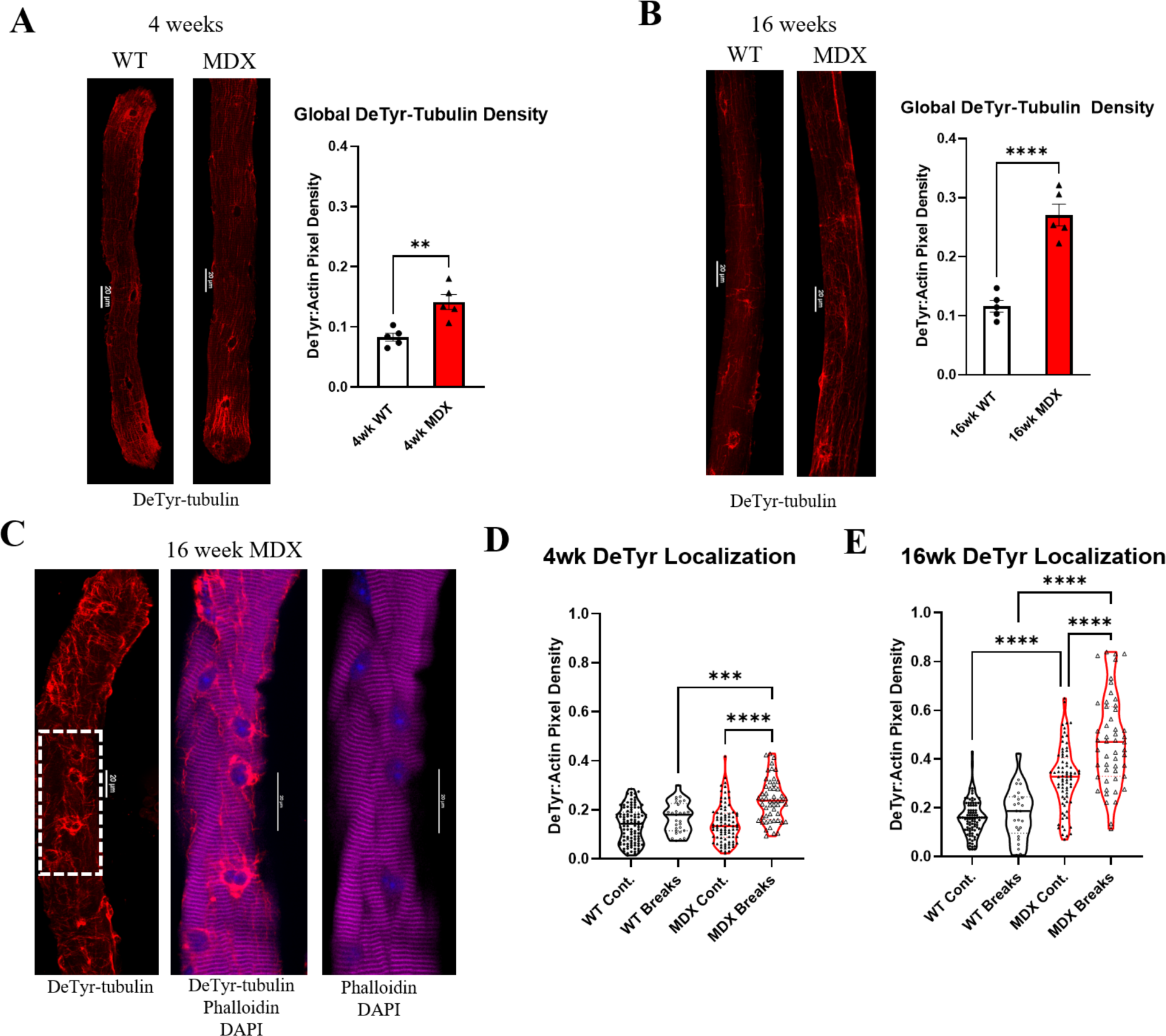
Detyrosination is enriched in dystrophic fibers and at the site of myofibrillar malformation. (A-B) We show the average deTyr density normalized to fiber size via the actin stain in fibers from wild-type and *mdx* mice 16 wks (n = 5 mice). Values are means ± SEM. Statistical significance determined using t-tests. **(C)** Representative images showing co- localization of detyrosinated tubulin with myofibrillar break sites. **(D-E)** Quantification of deTyr- tubulin density in regions with continuous striations across myofibrils vs. within break sites. Analysis was completed using one-way ANOVA with Šídák’s multiple comparisons test (**P < 0.01; ***P<0.001; ****P <0.0001).

In 16 wk WT muscle fibers we again find no difference in the density of deTyr -MTs within the regions of myofibrillar separation versus areas with otherwise normal myofibrillar connectivity. In the 16 wk *mdx* we again find a significant elevation in deTyr-MTs in areas of myofibrillar separation but now find these changes in areas of otherwise normal myofibrillar connectivity as well. (**Fig 7E**). Taken together, these results suggest that deTyr-MTs become abundant first between myofibrils then progress more globally throughout the myofibrillar structure as disease progresses.

### 3.6 Overexpression of VASH2-SVBP models the increased tubulin detyrosination, cytoskeletal stiffness, and myofibrillar malformations established as pathognomonic in *mdx*

Detyrosination is the reversible enzymatic cleavage of the COOH-terminal tyrosine from α- tubulin by vasohibin 1 (VASH1) or vasohibin 2 (VASH2) and their partner the small vasohibin binding protein (SVBP)^13, 38^. Examining the transcripts of these proteins at 4 wks (**Fig 8A-D**) finds no significant change in VASH1 yet a significant increase in VASH2 and SVBP in the *mdx*. At 16 wks the VASH1 remained unchanged, while VASH2 remained elevated in the *mdx*, and SVBP now exhibited no difference between genotypes (**Fig 8F-I**).

**Fig 8.**
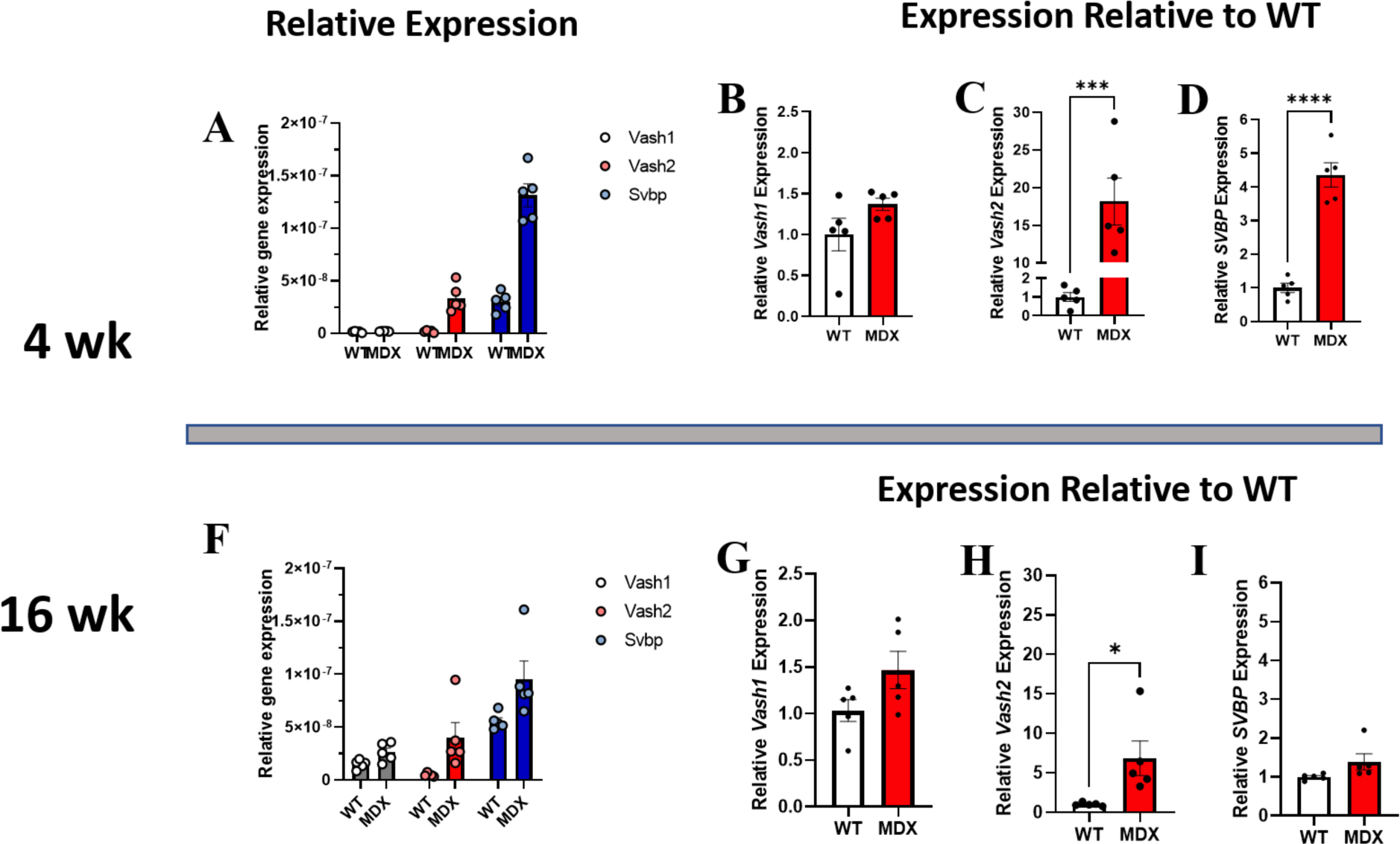
Expression of vasohibins and SVBP are increased in dystrophy. (A-D) Relative expression of VASH1, VASH2, and SVBP at 4 wks reveals **(B)** VASH1 expression is not significantly increased at 4 wks whereas **(C)** VASH2 expression is increased approximately 18- fold on average in the *mdx* and **(D)** and SVBP is increased approximately 4-fold. **(F)** Relative gene expression of both tubulin carboxypeptidases, and especially VASH1, appears increased in wild-type 16 wk compared to 4 wk and *mdx* expression relative to wild-type remains increased at 16 wks. **(G)** VASH1 shows a trend toward increased expression in *mdx*, only **(H)** VASH2 expression remains significantly increased, while its binding partner **(I)** SVBP is not significantly increased in *mdx* at 16 wks. All analysis was performed using one-way ANOVA with Šídák’s multiple comparisons test (*P < 0.05; ***P<0.001; ****P <0.0001).

Microtubules are essential for myofibrillar growth, maintenance and repair^21–24^. Given that bundles of deTyr modified MTs are associated with myofibrillar separations in 4 wk *mdx* muscle fibers that then progress to more significant changes at 16 wks, we sought to determine if an experimental increase in deTyr-MTs in 4 wk WT fibers was sufficient to recapitulate the changes seen in the 16 wk *mdx*. To this end we used electroporation to introduce VASH2-GFP/SVBP cDNA, or eGFP cDNA as a control, into the 4 wk old mouse FDB and examined the muscle fiber properties 5-7 days later (**Fig 9A-C**).

**Fig 9.**
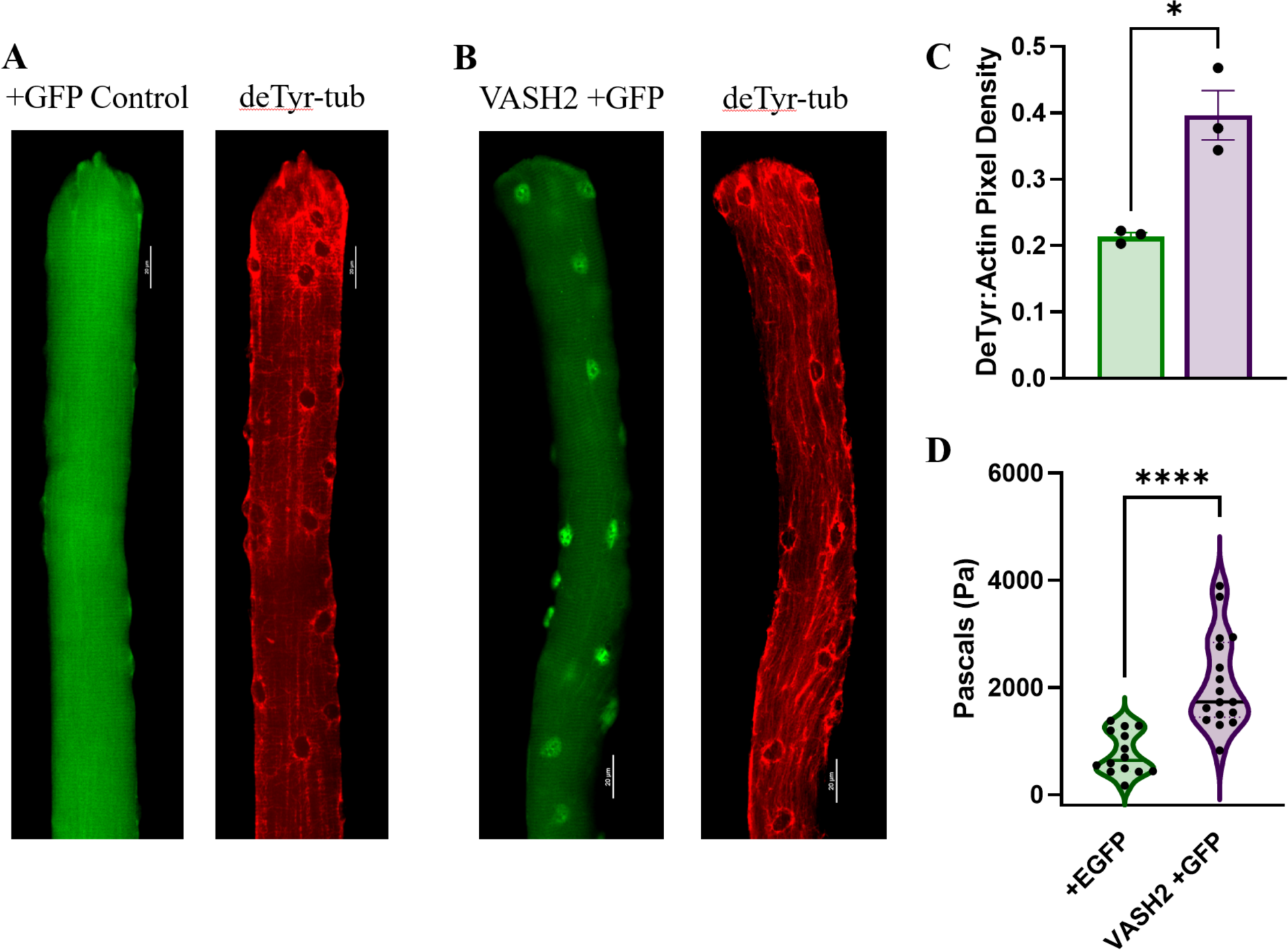
VASH2 overexpression increases the density of deTyr-MTs and the passive stiffness of the muscle fiber. **(A)** Representative image muscle fiber expressing EGFP(+) as a control finds deTyr-tubulin enriched MTs dispersed throughout the fiber. **(B)** VASH2+GFP expressing muscle fiber exhibits a denser and more bundled network of deTyr-tubulin enriched MTs. **(C)** Quantification of the density of deTyr-MTs finds a significant elevation with VASH2 overexpression (n=3 mice per condition; 10-15 fibers per mouse). **(D)** Nano-indentation (5µm/sec) measure of viscoelastic resistance finds increased stiffness in VASH2-GFP muscle fibers. (t-test, * P < 0.05; ****P < 0.0001).

In WT FDB fibers transduced with VASH2-GFP/SVBP we find an increased abundance of deTyr-enriched MT bundles when compared to the eGFP controls (**Fig 9A-C**). In *mdx* muscle fibers we previously linked the elevated levels of deTyr-MTs to an increase in cytoskeletal mechanics (i.e., stiffness)^17, 39^. Using nanoindentation to measure the viscoelastic properties we show a significant increase in passive stiffness inVASH2-GFP/SVBP over expressing FDB muscle fibers compared to the eGFP control (**Fig 9D**).

Our evidence suggests a link between the increased abundance of deTyr-enriched MTs and the altered myofibrillar structure in the 16 wk *mdx*. Consistent with this finding in *mdx* was a significant reduction in striation continuity in WT fibers overexpressing VASH2-GFP/SVBP determined by Z-line detection reporting (**Fig 10C**). Further evidence supporting a link between deTyr-MTs and myofibrillar malformations came from visual inspection that revealed a 21.3% occurrence of braided myofibrils in the VASH2-GFP/SVBP over-expressing myofibers with no evidence for this occurrence in controls. Concordant with our finding deTyr-enriched MTs co-registered with myofibrillar malformations in the *mdx* (**Fig7**), in VASH2-GFP/SVBP over- expressing myofibers we found bundled deTyr-MTs abundant only in areas with altered myofibrillar structure (**Fig 10E**).

**Fig 10.**
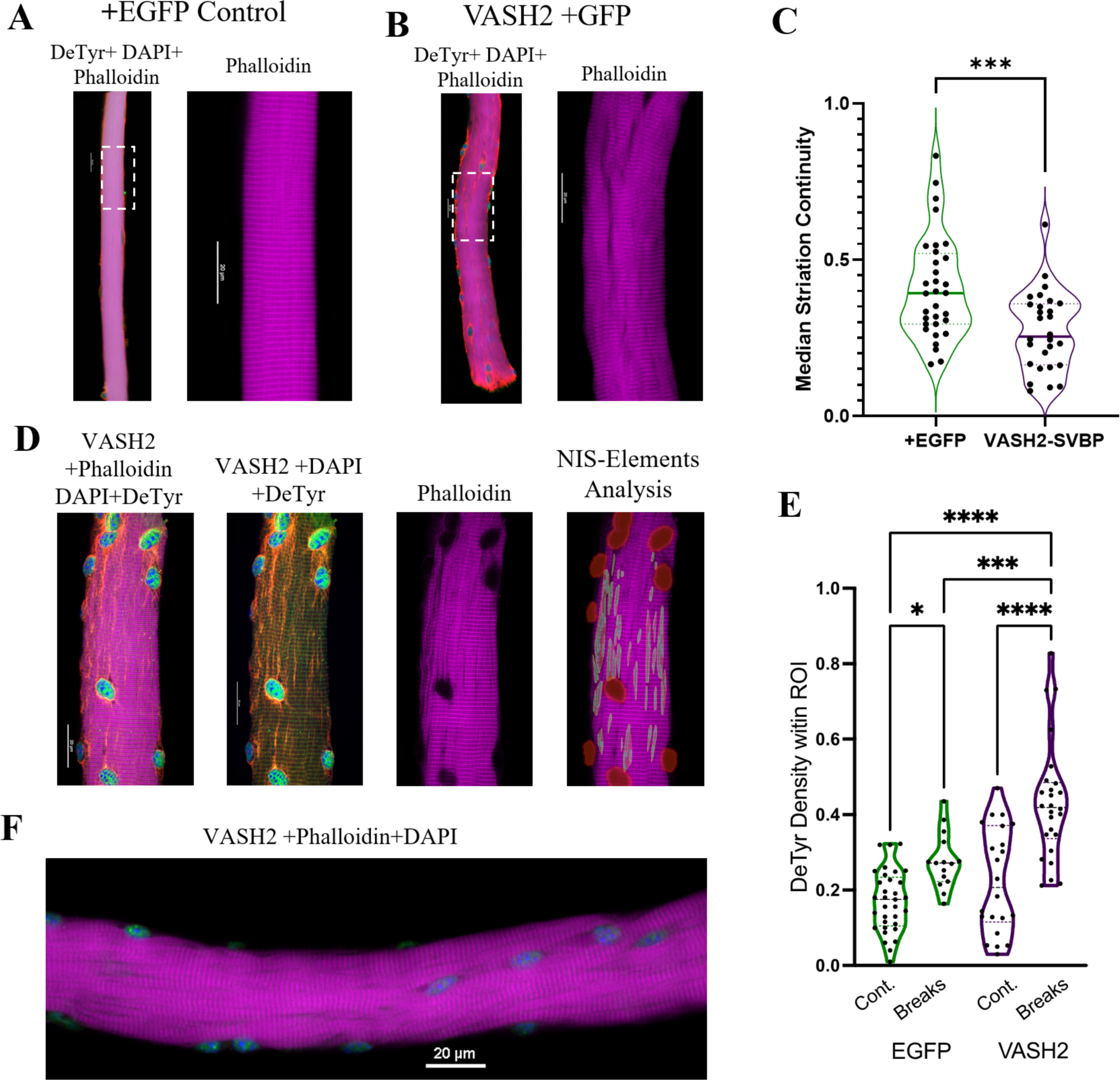
Detyrosinated microtubules are enriched at areas of Z-line disruption (A-B) Representative images of EGFP and VASH-SVBP electroporated fibers respectively, with insets showing the disparate morphology. A closer look at the inset **(B)** reveals extensive veneering of striations, perhaps indicative of the onset of myofibrillar “braiding”. **(C)** Quantification of striation continuity using Z-line detection. There was a statistically significant difference between groups as determined by t-test **(D)** Representative image of EGFP(+)VASH2 myofiber showing myofibrillar break sites identified as regions-of-interest (ROI) within NIS-Elements. **(E)** The density of the deTyr-MTs within each ROI was calculated and normalized to the respective area. There was a statistically significant difference between groups as determined by one-way ANOVA with Šídák’s multiple comparisons test (*F*(3,92) = 25.12). **(F)** Representative image of VASH2-SVBP electroporated FDB fiber with evidence of altered myofibrillar directionality (i.e., braiding). (* P < 0.05; ***P < 0.001; ****P < 0.0001).

Taken together, our results demonstrate that VASH2-GFP/SVBP overexpression in WT muscle is sufficient to model the altered myofibrillar structure that arises in the 16 wk *mdx*. We posit that disease altered microtubules are an early event in dystrophic pathology that predispose the altered myofibrillar structure in dystrophinopathies.

## Discussion

Altered myofibrillar structure is a consequence of dystrophic pathology in humans^40, 41^ and rodents ^3, 4, 19, 42^ that decreases force production and increases susceptibility to contraction injury. In fact, evidence from optical predictions^8^ and mathematical models^6^ suggests myofibril misalignment and myofibrillar stiffness are the dominant factors contributing to decreased isometric force and contraction velocity^6^ in dystrophinopathies. While the consequences of altered myofibrillar structure are well defined, the mechanisms that underlie their occurrence have been elusive.

Microtubules play a critical role in the myofibrillar growth, maintenance, and repair of striated muscle through the patterned recruitment of myosin, actin, mRNAs, and ribosomes for the assembly of sarcomeres^24^. Consistent with tubulin PTM’s as regulators of MT function, MTs enriched in deTyr-tubulin have been implicated in the regulation of mechanotransduction dependent ROS and Ca^2+^ signals, in the directional transport of cargo (i.e., lysosomes), and in the highly orchestrated myofibrillar assembly during myogenesis^25, 26^.

Here we show that myofibrillar malformations are not inherent to dystrophin’s absence, rather they arise in the *mdx* between 4 and 16 weeks of age coincident with the densification of deTyr-enriched MTs in these malformed areas. Transcriptional evidence of increased VASH2 and SVBP in the 4 wk *mdx* suggested deTyr-enriched MTs may be responsible for the myofibrillar alterations arising by 16 wks. Evidence supporting a causative link was established by showing that VASH2-GFP/SVBP overexpression in WT muscle fibers was sufficient to recapitulate the densification of deTyr-enriched MTs and the altered myofibrillar structure seen in the 16 wk *mdx*. These results suggest that the enhanced expression/activation of VASH2/SVBP in dystrophic muscle fibers is an early event in dystrophic pathology that drives the densification of deTyr-MTs to disrupt myofibrillar maintenance and/or repair.

This report extends our previous discovery implicating deTyr enriched MTs as an early event in DMD pathology that drives the excess mechanotransduction elicited Nox2-ROS and Ca^2+^ signals linked to dystrophic progression^16,17.18^. Together these results give strong support for disease altered MTs as negative disease modifiers early in DMD pathology. Given the transcriptional and proteomic evidence for these alterations in DMD patients^16, 43^, targeted therapeutics to reduce deTyr-MTs may be a viable option to slow dystrophic progression. As pharmacologic approaches are developed, future studies to genetically reduce VASH1, VASH2, or SVBP in *mdx* fibers will further advance our mechanistic understanding.

While here we focused on DMD, it is notable that myofibrillar malformations were modeled in WT muscle fibers by VASH2/SVBP overexpression. This result demonstrates that dystrophin’s absence is not obligate in this process, nor is the dysregulated signaling linked to dystrophic pathology. It is then tempting to speculate that the occurrence of myofibrillar malformations seen in aging muscle^44, 45^, in disparate genetic diseases^46^, and in conditions of supraphysiologic muscle growth^47, 48^ may be driven by this same axis. In this regard, future work profiling these conditions, and mechanisms that increase VASH/SVBP expression and activity, will likely yield insights of broad importance.

---

**Supplementary table 1.**
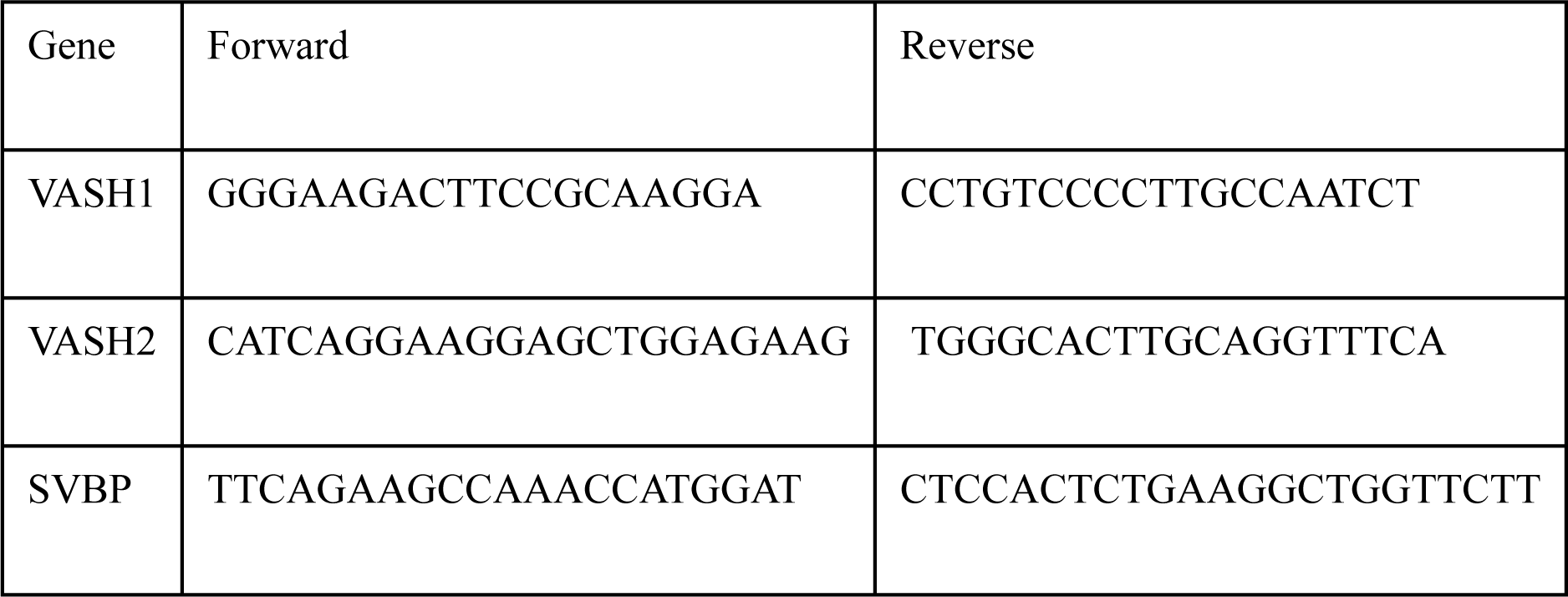
Primer sequences for RT-qPCR

## Notes

### Competing Interest Statement

The authors have declared no competing interest.

## References

1. Willingham, T. B., Kim, Y., Lindberg, E., Bleck, C. K. E. & Glancy, B. The unified myofibrillar matrix for force generation in muscle. Nat. Commun. 11, 3722 (2020).

2. Head, S. I., Williams, D. A., Stephenson, D. G. & Gage, P. W. Abnormalities in structure and function of limb skeletal muscle fibres of dystrophic mdx mice. Proc. R. Soc. Lond. B Biol. Sci. 248, 163–169 (1997).

3. Buttgereit, A., Weber, C., Garbe, C. S. & Friedrich, O. From chaos to split-ups - SHG microscopy reveals a specific remodelling mechanism in ageing dystrophic muscle. J. Pathol. 229, 477–485 (2013).

4. Liu, W., Raben, N. & Ralston, E. Quantitative evaluation of skeletal muscle defects in second harmonic generation images. J. Biomed. Opt. 18, 026005–026005 (2013).

5. Ding, J., Cong, Y. F., Liu, B., Miao, J. & Wang, L. Aberrant Protein Turn-Over Associated With Myofibrillar Disorganization in FHL1 Knockout Mice. Front. Genet. 9, (2018).

6. Stefanati, M., Torrente, Y. & Rodriguez Matas, J. F. Effect of myofibril architecture on the active contraction of dystrophic muscle. A mathematical model. J. Mech. Behav. Biomed. Mater. 114, 104214 (2021).

7. Ritter, P. et al. Myofibrillar Lattice Remodeling Is a Structural Cytoskeletal Predictor of Diaphragm Muscle Weakness in a Fibrotic mdx (mdx Cmah−/−) Model. Int. J. Mol. Sci. 23, 10841 (2022).

8. Schneidereit, D. et al. Optical prediction of single muscle fiber force production using a combined biomechatronics and second harmonic generation imaging approach. Light Sci. Appl. 7, (2018).

9. Liu, W. & Ralston, E. A new directionality tool for assessing microtubule pattern alterations. Cytoskeleton 71, 230–240 (2014).

10. Oddoux, S. et al. Misplaced Golgi Elements Produce Randomly Oriented Microtubules and Aberrant Cortical Arrays of Microtubules in Dystrophic Skeletal Muscle Fibers. Front. Cell Dev. Biol. 7, 1–18 (2019).

11. Prins, K. W. et al. Dystrophin is a microtubule-associated protein. J. Cell Biol. 186, 363–369 (2009).

12. Peris, L. et al. Motor-dependent microtubule disassembly driven by tubulin tyrosination. J. Cell Biol. 185, 1159–1166 (2009).

13. Aillaud, C. et al. Vasohibins/SVBP are tubulin carboxypeptidases (TCPs) that regulate neuron differentiation. Science 1453, 1448–1453 (2017).

14. Nieuwenhuis, J. et al. Vasohibins encode tubulin detyrosinating activity. Science 358, 1453– 1456 (2017).

15. Salomon, A. K. et al. Desmin intermediate filaments and tubulin detyrosination stabilize growing microtubules in the cardiomyocyte. Basic Res. Cardiol. 117, 53 (2022).

16. Khairallah, R. J. et al. Microtubules underlie dysfunction in duchenne muscular dystrophy. Sci. Signal. 5, ra56–ra56 (2012).

17. Kerr, J. P. et al. Detyrosinated microtubules modulate mechanotransduction in heart and skeletal muscle. Nat. Commun. 6, 1–14 (2015).

18. Prosser, B. L., Khairallah, R. J., Ziman, A. P., Ward, C. W. & Lederer, W. J. X-ROS signaling in the heart and skeletal muscle: stretch-dependent local ROS regulates [Ca^2+^]i. J. Mol. Cell. Cardiol. 58, 172–81 (2013).

19. Lovering, R. M. et al. Physiology, structure, and susceptibility to injury of skeletal muscle in mice lacking keratin 19-based and desmin-based intermediate filaments. Am. J. Physiol. - Cell Physiol. 300, C803–C813 (2011).

20. Head, S. I. Branched fibres in old dystrophic mdx muscle are associated with mechanical weakening of the sarcolemma, abnormal Ca2+ transients and a breakdown of Ca2+ homeostasis during fatigue. Exp Physiol 95, 641–56 (2010).

21. Pizon, V., Gerbal, F., Diaz, C. C. & Karsenti, E. Microtubule-dependent transport and organization of sarcomeric myosin during skeletal muscle differentiation. EMBO J. 24, 3781– 3792 (2005).

22. Scholz, D. et al. Microtubule-dependent distribution of mRNA in adult cardiocytes. Am. J. Physiol.-Heart Circ. Physiol. 294, H1135–H1144 (2008).

23. Denes, L. T., Kelley, C. P. & Wang, E. T. Microtubule-based transport is essential to distribute RNA and nascent protein in skeletal muscle. Nat. Commun. 12, 6079 (2021).

24. Dhanyasi, N., VijayRaghavan, K., Shilo, B.-Z. & Schejter, E. D. Microtubules provide guidance cues for myofibril and sarcomere assembly and growth. Dev. Dyn. 250, 60–73 (2021).

25. Gundersen, G. G., Khawaja, S. & Bulinski, J. C. Generation of a stable, posttranslationally modified microtubule array is an early event in myogenic differentiation. J. Cell Biol. 109, 2275–2288 (1989).

26. Chang, W. et al. Alteration of the C-terminal amino acid of tubulin specifically inhibits myogenic differentiation. J Biol Chem 277, 30690–8 (2002).

27. Aillaud, C. et al. Vasohibins/SVBP are tubulin carboxypeptidases (TCPs) that regulate neuron differentiation. Science 358, 1448–1453 (2017).

28. Nelson, D. M. et al. Variable rescue of microtubule and physiological phenotypes in mdx muscle expressing different miniaturized dystrophins. Hum. Mol. Genet. 27, 2090–2100 (2018).

29. Nelson, D. M. et al. Rapid, redox-mediated mechanical susceptibility of the cortical microtubule lattice in skeletal muscle. Redox Biol. 37, 101730 (2020).

30. Loehr, J. A. et al. NADPH oxidase mediates microtubule alterations and diaphragm dysfunction in dystrophic mice. eLife 7, 1–19 (2018).

31. Iyer, S. R. et al. Altered nuclear dynamics in MDX myofibers. J. Appl. Physiol. 122, 470–481 (2017).

32. McHugh, M. P. & Tyler, T. F. Muscle strain injury vs muscle damage: Two mutually exclusive clinical entities. *Transl*. SPORTS Med. 2, 102–108 (2019).

33. Goodall, M. H., Ward, C. W., Pratt, S. J. P., Bloch, R. J. & Lovering, R. M. Structural and functional evaluation of branched myofibers lacking intermediate filaments. Am. J. Physiol.- Cell Physiol. 303, C224–C232 (2012).

34. Lovering, R. M., Michaelson, L. & Ward, C. W. Malformed mdx myofibers have normal cytoskeletal architecture yet altered EC coupling and stress-induced Ca2+ signaling. Am. J. Physiol. Cell Physiol. 297, C571–80 (2009).

35. Morris, T. A. Computational and Image Analysis Techniques for Quantitative Evaluation of Striated Muscle Tissue Architecture. (University of California, Irvine, 2021).

36. Otsu, N. A Threshold Selection Method from Gray-Level Histograms. IEEE Trans. Syst. Man Cybern. 9, 62–66 (1979).

37. Morris, T. A. et al. Striated myocyte structural integrity: Automated analysis of sarcomeric z-discs. PLOS Comput. Biol. 16, e1007676 (2020).

38. Ramirez-Rios, S. et al. VASH1-SVBP and VASH2-SVBP generate different detyrosination profiles on microtubules. 2022.06.02.494516 Preprint at https://doi.org/10.1101/2022.06.02.494516 (2022).

39. Coleman, A. K., Joca, H. C., Shi, G., Lederer, W. J. & Ward, C. W. Tubulin acetylation increases cytoskeletal stiffness to regulate mechanotransduction in striated muscle. J. Gen. Physiol. 153, e202012743 (2021).

40. Murach, K. A. et al. Starring or Supporting Role? Satellite Cells and Skeletal Muscle Fiber Size Regulation. Physiology 33, 26–38 (2017).

41. Olivé, M. et al. Desmin-related myopathy: Clinical, electrophysiological, radiological, neuropathological and genetic studies. J. Neurol. Sci. 219, 125–137 (2004).

42. Kiriaev, L. et al. Lifespan Analysis of Dystrophic mdx Fast-Twitch Muscle Morphology and Its Impact on Contractile Function. Front. Physiol. 12, (2021).

43. Capitanio, D. et al. Comparative proteomic analyses of Duchenne muscular dystrophy and Becker muscular dystrophy muscles: changes contributing to preserve muscle function in Becker muscular dystrophy patients. J. Cachexia Sarcopenia Muscle 11, 547–563 (2020).

44. Pichavant, C. & Pavlath, G. K. Incidence and severity of myofiber branching with regeneration and aging. Skelet Muscle 4, 9 (2014).

45. Grounds, M. D. Therapies for sarcopenia and regeneration of old skeletal muscles: more a case of old tissue architecture than old stem cells. Bioarchitecture 4, 81–7 (2014).

46. Seto, J. T. et al. Deficiency of α-actinin-3 is associated with increased susceptibility to contraction-induced damage and skeletal muscle remodeling. Hum. Mol. Genet. 20, 2914– 2927 (2011).

47. Antonio, J. & Gonyea, W. J. Muscle fiber splitting in stretch-enlarged avian muscle. Med. Sci. Sports Exerc. 26, 973–977 (1994).

48. Murach, K. A., Dungan, C. M., Peterson, C. A. & McCarthy, J. J. Muscle Fiber Splitting Is a Physiological Response to Extreme Loading in Animals. Exerc. Sport Sci. Rev. 47, 108–115 (2019).

